# Pervasive ancestry bias in variant effect predictors

**DOI:** 10.1101/2024.05.20.594987

**Authors:** Ankit K. Pathak, Nikita Bora, Mihaly Badonyi, Benjamin J. Livesey, SG10K_Health Consortium, Joanne Ngeow, Joseph A. Marsh

**Affiliations:** MRC Human Genetics Unit, Institute of Genetics and Cancer, University of Edinburgh, Edinburgh, UK; Lee Kong Chian School of Medicine, Nanyang Technological University, Singapore; Oncology Academic Clinical Program, Duke-NUS Medical School, Singapore; Cancer Genetics Service, Division of Medical Oncology, National Cancer Centre Singapore, Singapore

## Abstract

Variant effect predictors (VEPs) – computational tools that assess the potential impact of genetic variants – have become increasingly vital for clinical variant interpretation. Currently, most VEPs used in variant prioritisation have been trained on datasets of clinically curated or population-derived variants. These datasets, however, disproportionately represent individuals of European descent. We hypothesised that this bias may lead to unequal VEP performance across different populations. To test this, we evaluated the scoring patterns of 52 VEPs for missense variants across 14 ancestry groups. We observe striking disparities: some VEPs predict a markedly higher proportion of damaging variants in underrepresented populations, such as those of Malay descent, compared to individuals of European ancestry. In contrast, VEPs that do not rely on clinical or population data predict more consistent pathogenicity burdens across ancestry groups. Moreover, we could closely link these discrepancies across methods to biases in training data. Our findings underscore the urgent need to adopt tools that minimise ancestry bias to ensure fairer and more accurate variant effect prediction and genetic diagnoses for all populations.

## INTRODUCTION

Over the past two decades, rapid advances in population sequencing technologies have produced a vast catalogue of human genomic variants, greatly enhancing our understanding of genetic diseases and transforming clinical care. Yet, these benefits have not been equitable. Genomic studies have historically prioritised individuals of European descent, resulting in poor representation of other ancestry groups^1^. Despite efforts to diversify cohorts, sequences and variants of European ancestry continue to dominate genomic databases^2^. This imbalance has profound consequences, increasing the risk of misdiagnosis for underrepresented populations^3,4^. Addressing these systemic biases through ongoing evaluation and strategic solutions is critical for achieving equitable healthcare outcomes for all.

Variant effect predictors (VEPs) are computational tools designed to assess the phenotypic impacts of genetic variants, particularly their potential to cause disease. Most research has targeted missense variants – single amino acid substitutions – since they dominate known variants of uncertain significance^5^. Over time, a growing number of VEPs have emerged, steadily improving in their ability to identify damaging missense variants^6–8^. Although current guidelines recommend using VEPs only as *supporting* evidence in making genetic diagnosis^9^, they are widely employed in clinical variant prioritisation, with ongoing efforts to increase their diagnostic weight^10^.

Most VEPs currently utilised in clinical practice are based on supervised machine learning models, trained on datasets of variants that have been clinically classified as either pathogenic or benign. In addition, these predictors also often include information about population allele frequency. We refer to these as *clinical-trained* VEPs according to our previously described classification scheme^6^. Despite their widespread use, these predictors are inherently associated with concerns regarding data circularity – a phenomenon where training and evaluation data overlap – leading to potential bias and inflated performance estimates^11^. This can arise from variant-level (type 1) circularity, where the same variants are used in both training and performance evaluation, or gene-level (type 2) circularity, where variants from the same or homologous genes contribute to both training and assessment. Given this, we hypothesised that clinical-trained VEPs could be particularly vulnerable to ancestry representation bias (hereafter referred to as ancestry bias) stemming from the composition of the variants on which they are trained.

In contrast, *population-tuned* VEPs do not rely on clinical variant classifications for training, but instead incorporate human population variants to inform their models. This can be achieved in several ways: by incorporating population variants directly into the model, using them to calibrate predictions across different genes, or fine-tuning the model with allele frequency data. This strategy has been growing in popularity, with notable recent examples including AlphaMissense^12^ and popEVE^13^. Theoretically, these predictors should be less prone to data circularity than clinical-trained VEPs, a notion supported by recent performance evaluations^6^. However, the risk is not entirely eliminated; for instance, allele frequencies, which are integral to clinical genomic interpretation^9^, can introduce circularity when these methods are evaluated on clinical benchmarks^14^. Moreover, given their exposure to human population variants, these models could still be susceptible to ancestry bias.

Finally, a third category of predictors, the *population-free* VEPs, are entirely unexposed to clinical or human population variants. Instead, these models primarily leverage evolutionary information from multiple sequence alignments or protein language models^15^, and increasingly on protein structural features^16^. This class largely overlaps with what is often referred to as “unsupervised” models, though it can encompass supervised models trained on alternative data sources, such as deep mutational scanning datasets. In recent years, several new population-free VEPs have shown remarkable performance, often surpassing clinical-trained methods in terms of correspondence with functional assays and identification of pathogenic variants^6,17^. However, despite this emerging potential, these tools remain underutilised in clinical genomic interpretation, likely because they are not explicitly designed to predict pathogenicity. Crucially, these population-free VEPs should be completely free of both data circularity or ancestry bias, providing a potential strategy for fairer variant interpretation across diverse populations.

## RESULTS

In this study, we investigated how ancestry bias in the genetic variants used for training variant effect predictors influences their performance. Specifically, we hypothesised that clinical-trained and population-tuned VEPs might exhibit differential behaviour across ancestry groups, reflecting the likely skewed representation of these groups in their training data. To test this, we first compile sets of missense variants from 14 ancestry groups defined in three large-scale population sequencing studies. From gnomAD v4.1^18^, we include African/African American (AFR), Admixed American (AMR), Ashkenazi Jewish (ASJ), East Asian (EAS), Finnish (FIN), Middle Eastern (MID), non-Finnish European (NFE) and South Asian (SAS). From the Mexico City Prospective Study (MCPS)^19^, we include African from Mexico City (AMX), European from Mexico City (EMX) and Indigenous Mexican (IMX). From Singapore’s SG10K Health project^20^, we include Chinese (CHI), Indian (IND), and Malay (MAL).

Importantly, we acknowledge that discrete ancestry classifications have inherent limitations, as ancestry is a continuous trait not fully captured by such categorical labels^21^. For this analysis, we adopt the 14 ancestry groups as defined by the respective sequencing studies, not to assert their precision or completeness, but to examine how VEP performance varies when applied to variants from these diverse populations. Our hypothesis relies on the assumption that these ancestry groups differ in their representation in VEP training datasets, which could potentially lead to differential performance. Thus, the use of these categorical labels serves as a practical framework to explore potential biases in VEP performance across ancestry groups, rather than as a definitive representation of human genetic diversity.

To standardise comparisons between ancestry groups with varying sample sizes, we first filter for single nucleotide missense variants with allele frequencies between 0.1% and 1% (Figure 1A; see Methods). This range ensures that variants are detectable in all cohorts, including the smallest ancestry groups (for *e*.*g*., 1,608 individuals in Malay), while avoiding the overrepresentation of rare variants in larger cohorts, which would confound comparisons. Additionally, it excludes very common variants (>1%), which are more likely to be benign and less relevant to pathogenicity assessments.

**Figure 1.**
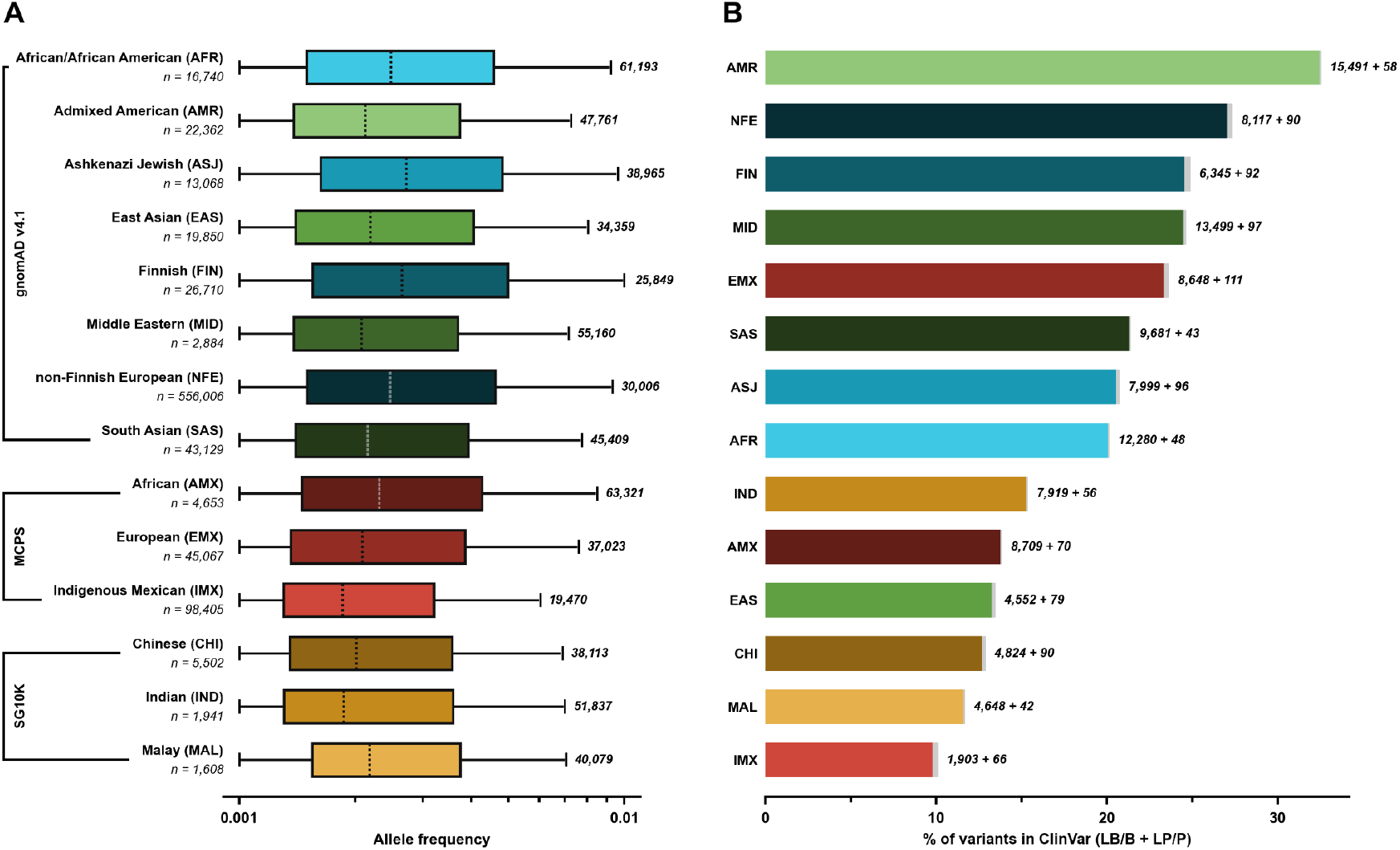
Ascertainment bias in variant knowledge across ancestry groups. **(A)** The box plot illustrates the distribution of allele frequencies for single nucleotide missense variants analysed in this study across 14 ancestry groups from three datasets: gnomAD v4.1, the Mexico City Prospective Study (MCPS), and Singapore’s SG10K Health project (SG10K). For each ancestry group, the sample size (n) is indicated on the left, while the number of variants examined is shown on the right. Only variants with allele frequencies between 0.1% and 1% were included to enable consistent comparisons across groups. This criterion helps avoid bias from ancestry groups with more extensive sequencing, which could otherwise introduce an over-representation of rare variants and complicate analyses. Outliers have been excluded for visual clarity. **(B)** The proportion of variants classified as either (likely) benign (coloured) or (likely) pathogenic (grey) in ClinVar is shown for each ancestry group, along with their corresponding counts. Differences in these proportions highlight the extent of biases in the ascertainment of variant knowledge across different ancestry groups.

In Figure 1B, we compare the proportion of these variants assigned clinical classifications in ClinVar^22^ across ancestry groups. As expected for population variants, far more have benign (LB/B) than pathogenic (LP/P) labels. However, striking differences emerge in classification rates across populations. The highest classification rates are observed in the non-Finnish European (NFE, 27.4%) and Admixed American (AMR, 32.6%) groups, whereas variants from Malay (MAL, 11.7%) and Indigenous Mexican (IMX, 10.1%) groups are much less likely to have labels. These disparities highlight the significant ascertainment bias in variant knowledge across ancestry groups, where variants in individuals in certain populations have a substantially higher likelihood of being clinically interpreted^23^.

To compare the scoring patterns of different VEPs across ancestry groups, variant effect scores from each predictor were rank-normalised to a scale of 0 (least likely to be pathogenic) to 1 (most likely to be pathogenic). This normalisation was essential to ensure comparability, as different predictors employ distinct scoring scales. Figure 2A depicts the distribution of these rank scores for several representative VEPs across all 14 ancestry groups. First, we examine the population-free models CPT-1^24^, ESM-1b^25^, and GEMME^26^, which, by design, should be unaffected by ancestry bias and therefore reflect only genuine differences between ancestry groups. All three VEPs show largely consistent patterns, with the highest proportion of damaging variants observed in IMX, FIN and ASJ, and the lowest proportion in AMR and AFR. However, the overall distributions are relatively flat, with only modest differences between ancestry groups. For instance, when considering variants in the top quartile (scores >0.75) as damaging, CPT-1 predicts the fewest damaging variants in AMR (22.3%) and the most in IMX (29.0%). Similar ranges are observed for ESM-1b (22.3-29.2%) and GEMME (22.7-28.4%) for ancestry groups with the fewest and the most damaging variants.

**Figure 2.**
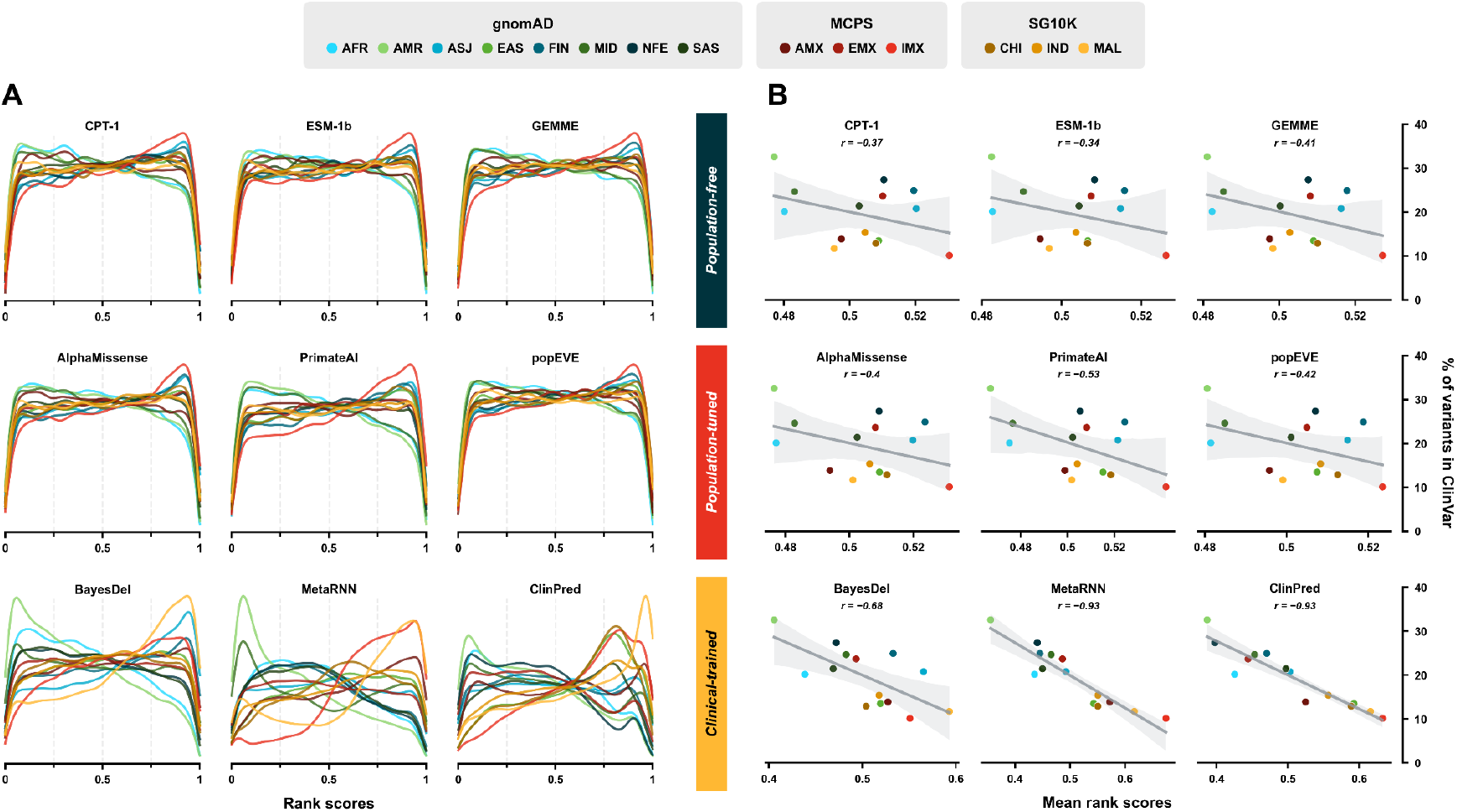
Variant annotation disparities significantly influence scoring patterns of clinical-trained VEPs. **(A)** The density plots depict the distribution of rank scores for nine VEPs across the 14 ancestry groups. Three representative VEPs are shown from each of the three classes based on their training method: *population-free, population-tuned*, and *clinical-trained*. The score gradient ranges from predicted benign (lower scores) to predicted pathogenic (higher scores) variants. **(B)** For the same set of VEPs, the correlation between the mean rank scores across the 14 ancestry groups and the percentage of these variants represented in ClinVar is displayed. The line indicates the best linear regression model fit, with the shaded region representing the 95% confidence interval. The Pearson correlation coefficient (*r*) quantifies the strength of this relationship.

Next, we analysed three population-tuned VEPs. Interestingly, we observed slightly larger ranges of predicted damaging (top quartile) variants for AlphaMissense (21.7-29.6%) and PrimateAI^27^ (20.9-30.8%), but not popEVE (22.5-28.3%), when comparing ancestry groups with the fewest and the most damaging variants. Broadly speaking, however, the distributions align closely with those of population-free VEPs, suggesting that these models are not strongly biased by their exposure to population variants.

In contrast to the population-free and population-tuned VEPs, the clinical-trained VEPs BayesDel^28^, MetaRNN^29^, and ClinPred^30^ show radical differences across ancestry groups, with heavily skewed score distributions. The range in the proportion of predicted damaging variants across ancestry groups is much larger for all three VEPs: BayesDel (15.1-38.3%), MetaRNN (11.5-45.5%) and ClinPred (11.3-41.6%). Notably, MAL – one of the least represented ancestry groups in ClinVar – was predicted by all three clinical-trained VEPs to have the highest pathogenicity burden, whereas the population-free and population-tuned VEPs rank it as intermediate compared to other ancestry groups. Furthermore, while there are small similarities in the distributions across ancestry groups, the three clinical-trained VEPs substantially vary amongst themselves, suggesting that each model is biased in distinct ways.

To better understand the origins of these disparate trends observed in clinical-trained VEPs, we calculated the mean rank scores for variants from each ancestry group using each VEP. This yielded a single value per group per VEP, with higher values indicating variants that are predicted to be more damaging overall. Figure 2B illustrates these mean rank scores against the proportion of variants present in ClinVar for each ancestry group, as shown in Figure 1B. Population-free and population-tuned VEPs showed relatively weak, non-significant Pearson correlation coefficients, ranging from -0.33 to -0.42, with PrimateAI displaying a slightly stronger correlation of -0.53. In contrast, clinical-trained VEPs showed much stronger correlations: -0.68 for BayesDel, and -0.93 for MetaRNN and ClinPred. Thus, the pathogenicity burden predicted by MetaRNN and ClinPred across ancestry groups can almost entirely be explained by the proportion of variants from those groups that have clinical classifications in ClinVar, suggesting that these VEPs are heavily biased by their training data.

To demonstrate the generality of these trends, we extended our analysis to 52 VEPs – 14 population-free, 6 population-tuned, and 32 clinical-trained. The score distributions and correlations with ClinVar classifications for all VEPs are shown in Figures S2 and S3. The heatmap in Figure 3 summarises the ancestry-wise trends for each VEP by illustrating their mean rank scores. The VEPs were then sorted by the standard deviation of their mean rank scores across ancestry groups. This arrangement highlighted the extent of variability predicted by VEPs between the ancestry groups, with higher standard deviations indicating greater predicted differences between groups.

**Figure 3.**
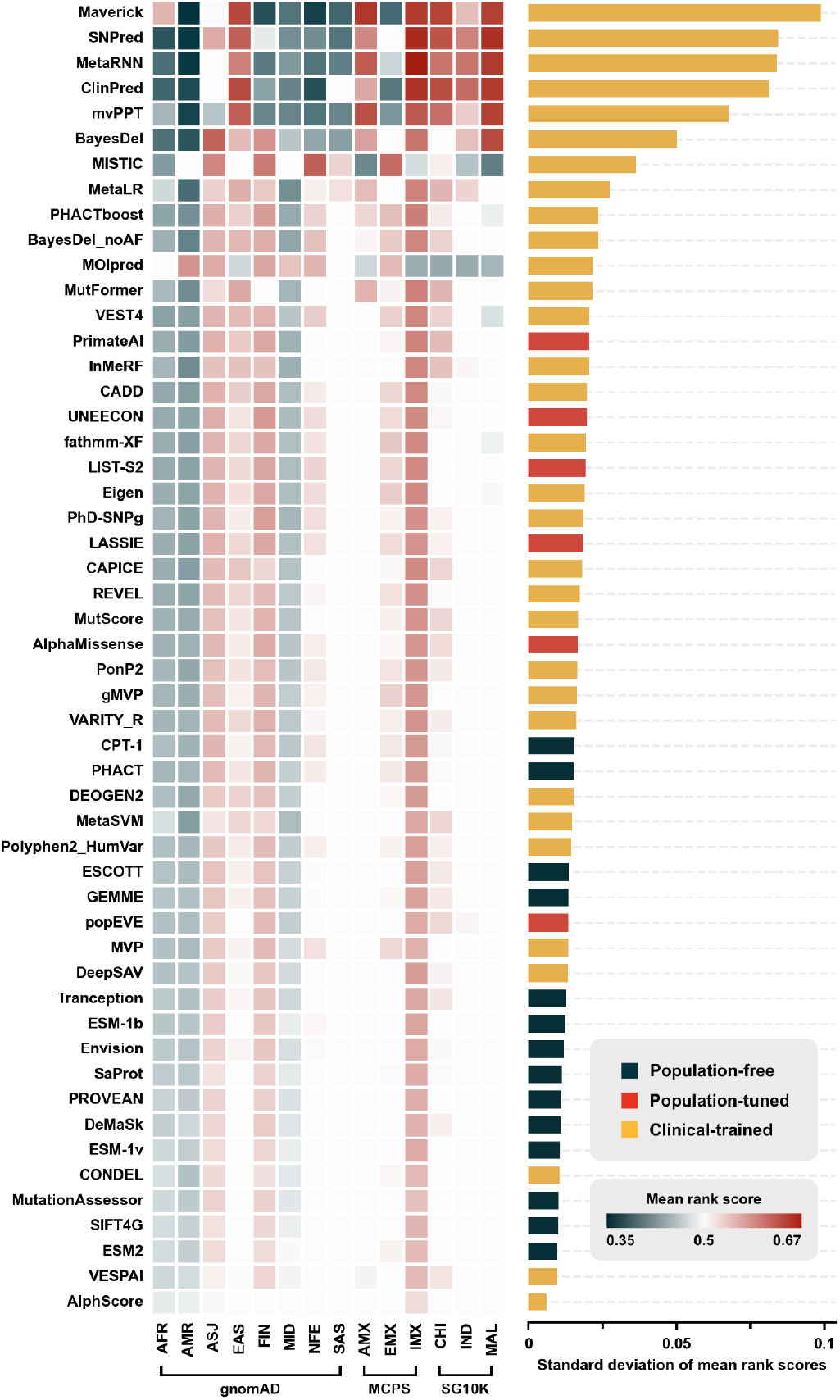
Population-free VEPs predict lower variability across ancestry groups compared to clinical-trained and population-tuned VEPs. The heatmap presents the mean rank scores of 52 VEPs across the 14 ancestry groups, with VEPs ordered by descending standard deviation of these scores, as shown in the accompanying bar plot. This arrangement highlights the influence of ancestry bias, with clinical-trained VEPs showing the largest deviations between ancestry groups, population-tuned VEPs displaying moderate, while population-free VEPs exhibit the least.

Strikingly, the 13 VEPs with the largest standard deviations were all clinical-trained. While the top seven appear to be clear outliers, showing extreme levels of population variability, most other clinical-trained and population-tuned VEPs still depict much larger standard deviations than population-free models. In fact, 24/32 clinical-trained and 5/6 population-tuned VEPs have standard deviations larger than all 14 population-free VEPs. These trends highlight the pervasiveness of ancestry bias on variant effect prediction, particularly in clinical-trained VEPs which appear to be highly susceptible to biases stemming from the ancestral compositions of their training datasets.

One possible argument for the results in Figure 3 is that the greater variability observed across ancestry groups by clinical-trained VEPs simply represents superior performance in identifying genuine differences. Indeed, if tested purely on their ability to discriminate between known pathogenic and benign variants, measured by the receiver operating characteristic (ROC) area under the curve (AUC), the six top-performing predictors also show the largest differences between ancestry groups (Figure 4). However, this analysis does not account for the issue of data circularity, as many of the pathogenic and benign variants will have been directly used in training^11^. Importantly, these six predictors also show very strong correlations between the mean VEP scores and the fraction of variants with classifications in ClinVar across ancestry groups (Figure S3), further suggesting overfitting to clinically annotated variants. Given that these annotated variants significantly overlap with those used for calculating ROC AUC, this overfitting likely inflates their apparent performance. Moreover, if the large differences between ancestry groups observed by these clinical-trained VEPs were solely to their superior predictive ability, similar patterns should appear across predictors. This is not the case, as shown in Figures 2 and 3, pointing instead to potential ancestry biases in these predictors.

**Figure 4.**
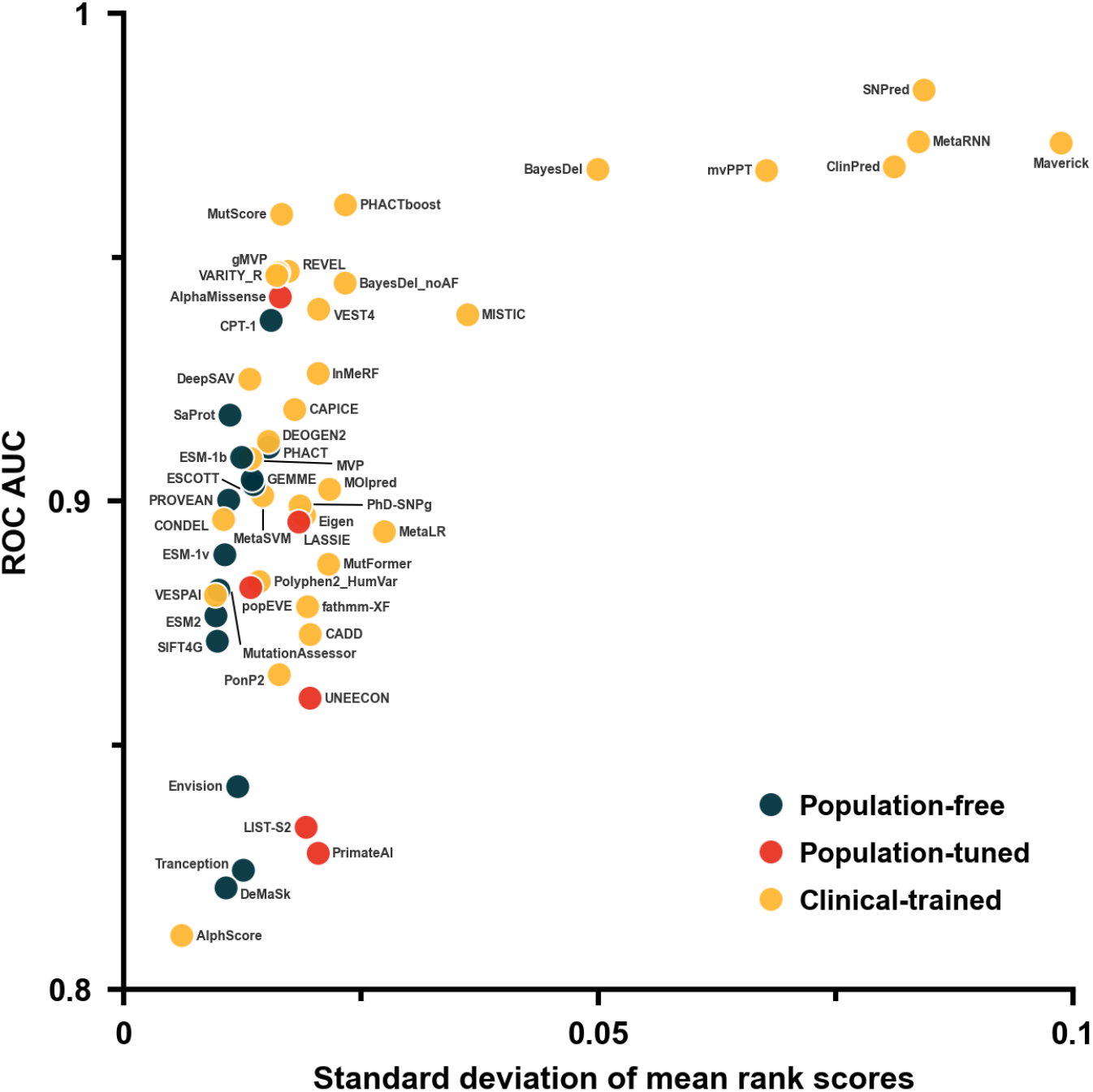
Overfitting to clinical variants drives ancestry bias in clinical-trained VEPs. The relationship between ROC AUC scores, for distinguishing between (likely) benign and (likely) pathogenic variants from ClinVar, and the standard deviation of mean rank scores from Figure 3 is plotted. Clinical-trained VEPs appear right-shifted, highlighting the impact of data circularity and ancestry bias in their training, particularly among the top-performing predictors.

Beyond these six most biased predictors, there is only a weak correspondence between the ROC AUC and the standard deviation of mean VEP scores. Some relationship is expected: there are genuine differences between ancestry groups as reflected by the consistent patterns observed in the lower part of Figure 4, particularly by population-free VEPs that are inherently free of population bias.. Thus, it is not surprising that CPT-1, the top-performing population-free VEP in pathogenicity prediction, also has the highest standard deviation. However, clinical-trained VEPs collectively exhibit a noticeable rightward shift in Figure 4, meaning they show larger differences between ancestry groups than population-free predictors with comparable pathogenicity prediction performance. This aligns with our recent findings that most clinical-trained VEPs perform relatively better in pathogenicity prediction than when evaluated against independent deep mutational scanning data, also highly suggestive of overfitting to clinical and population variants^6^.

## DISCUSSION

Efforts are underway to strengthen the clinical role of VEPs, particularly by elevating the evidence strength they can provide for genetic diagnosis^31,10,32^. This is important, as recent improvements in VEP performance strongly indicates that they can provide greater utility than their current *supporting* evidence status suggests. However, if VEPs are to be more widely adopted in clinical practice, it is imperative that their bias be minimised. In this study, we identified pervasive ancestry bias across most VEPs trained using clinical or population-derived variants. For instance, BayesDel, a tool frequently used by clinicians and recommended for providing *strong* evidence of pathogenicity^10,32^, exhibited some of the most pronounced ancestry bias in our analyses. This level of bias clearly renders it unsuitable for clinical variant interpretation. Other VEPs with established pathogenicity or benignity thresholds, such as REVEL^33^, VEST4^34^ and CADD^35^, showed moderate ancestry bias, though less pronounced than BayesDel, and should only be used in clinical contexts with great caution. It is also important to emphasise that the calibration of evidence strengths for VEP scores also relies on clinical and population variants^10,32^, meaning biases present in these datasets could also compromise the reliability of calibration.

The ancestry bias observed in this study stems from training datasets that reflect skewed ancestry compositions, driven by both clinical ascertainment biases and imbalanced population variant distributions. Variants observed in specific populations are more likely to have been investigated by clinicians, and therefore classified as pathogenic or benign^23^. However, biases in the population variant datasets used by VEPs may also play a significant role. For instance, some predictors heavily rely on allele frequency as a feature, which, as mentioned earlier, can inflate apparent performance metrics since allele frequency is often directly used in clinical variant classifications. This was highlighted in the recent Critical Assessment of Genome Interpretation (CAGI) Annotate-All-Missense challenge^8^, which demonstrated that the exceptional apparent performance of VEPs that directly utilise allele frequencies is almost entirely driven by common variants, with their performance dropping dramatically when applied to rare pathogenic and benign variants. Thus, it is informative to consider the case of BayesDel, as our analyses also included BayesDel_noAF, a version that does not utilise allele frequencies. While BayesDel_noAF shows lesser variability in scoring of variants from different ancestry groups compared to BayesDel, it still ranks 10th overall in terms of its standard deviation, suggesting that both allele frequency and training on clinical labels are important contributors to ancestry bias.

Not all clinical-trained VEPs show signs of ancestry bias. VARITY_R^36^, in particular, is notable for its strong performance in variant effect prediction, ranking as the top-performing clinical-trained model against functional assay data in our recent benchmark^6^, while showing minimal signs of ancestry bias in this study. This is likely attributable to its unique training strategy, which deliberately minimises circularity and bias by excluding features influenced by variant annotations and protein identity. This approach should serve as a model for future clinical-trained models. We also note that VARITY_R has been calibrated to provide *strong* evidence for variant pathogenicity assessment^32^, making it a good choice for clinical diagnosis if one wishes to use a clinical-trained model.

Population-tuned VEPs, on the other hand, tend to predict differences between ancestry groups that are intermediate to population-free and clinical-trained methods. For example, AlphaMissense displays a greater standard deviation across ancestry groups than any population-free model, suggesting a relatively modest but non-zero degree of ancestry bias. In contrast, popEVE closely resembles population-free VEPs, likely because it has not been tuned using any knowledge of specific variants, but instead utilises population variants for gene-level calibration of variant effect scores. This strategy should, in principle, avoid introducing ancestry bias, provided there are no significant differences between ancestry groups in the distribution of variants across genes. While the recent performance of population-tuned VEPs is promising, we recommend that, where feasible, developers of these methods also consider providing population-free scores alongside their population-tuned predictions to help assess potential biases.

One potential strategy to mitigate bias in clinical-trained and population-tuned VEPs is to base them on more representative populations. This would lessen disparities in VEP performance across ancestry groups, and we endorse this approach as more diverse clinical and population variant datasets emerge^37^. However, while there are many important reasons for increasing the diversity of genomic datasets^38–42^, this alone would not fully address the fundamental issue of many current VEPs being overfit to their training data, excelling with known variants but potentially failing to generalise to novel ones.

It is evident that certain population-free VEPs, like CPT-1, outperform most clinical-trained VEPs in pathogenicity prediction while not being prone to ancestry bias or circularity-related inflation of performance estimates. Notably, ESM-1b, a population-free VEP, has demonstrated the potential to achieve *strong* evidence strengths in pathogenic variant classification^32^, suggesting that others like CPT-1 would easily reach this level if calibrated. Thus, our major recommendation is that researchers and clinicians should more widely adopt population-free VEPs to ensure more robust and unbiased variant interpretation.

We note that population-free models could theoretically be affected by subtle ancestry bias, as they may be influenced by sequences that are close to their training data^43–45^. However, this mostly entails species-level effects and is unlikely to affect their ancestry-level predictions. In principle, if a reference sequence used by an unsupervised model represents a European variant that is different from other ancestry groups, this could introduce a level of bias even in population-free VEPs. However, given that we have only considered variants with allele frequencies <1% here, this should have no impact on our analyses. Furthermore, it should also be possible to reduce or eliminate this hypothetical bias by excluding human sequences from sequence alignments or protein language models, or using reference sequences that correspond to the population of interest.

Despite the strong pathogenicity prediction performance of recent population-free VEPs, it is possible that training on clinical or population-derived variants can improve predictions in ways unrelated to circularity. However, as we have extensively discussed^6^, it remains extremely challenging to assess this fairly. Therefore, while we recognise that many researchers and clinicians will prefer using clinical-trained models, particularly those they are familiar with, we strongly recommend incorporating population-free VEPs alongside other methods to help identify potential biases. For instance, if a clinical-trained VEP reaches a *strong* level of evidence for a variant, it is worth verifying whether a top population-free VEP (*e*.*g*. CPT-1) yields a similar result. If discrepancies arise, it is important to consider whether these could be linked to differences in variant prevalence across ancestry groups.

We do observe that there are clear, consistent tendencies seen by population-free and many other VEPs for certain ancestry groups to have variants that, on average, are either more or less damaging than others. These patterns likely reflect underlying population histories, for *e*.*g*., African populations tend to have greater genetic diversity^46^, while populations shaped by isolation and bottlenecks, such as the Finnish, are known to carry deleterious alleles at higher frequencies than non-bottlenecked populations, elevating their overall pathogenicity burden^47^. While these trends are interesting, we wish to emphasise that quantifying or explaining differences between ancestry groups is not the focus of our study.

Finally, beyond computational approaches, experimental methods also offer significant promise for reducing bias. High-throughput strategies for measuring variant effects at scale, known as multiplex assays of variant effect (MAVEs)^48^, have gained rapid popularity. Recent studies have demonstrated that MAVEs, when applied to genes such as BRCA1 and TP53, can reclassify variants of uncertain significance (VUS) at higher rates in non-European populations, helping address inherent disparities in traditional methods^49^. For now, however, the number of human genes with high-quality MAVEs remains limited^50^, and the lack of standardisation across different assays hampers the direct comparability of variant effect scores, limiting their broader clinical applicability. Thus, variant effect predictors will remain indispensable in clinical variant interpretation for the foreseeable future. The insights from our work underscore an urgent need to refine these predictive models, ensuring they are not only more accurate but also equitable across diverse populations. Addressing these biases is not just a technical challenge – it is a critical step toward more inclusive and just genomic medicine.

## METHODS

### Population Sequencing Datasets

We used three large-scale population sequencing datasets to gather genomic variants associated with numerous ancestry groups – gnomAD v4.1^18^, Mexico City Prospective Study (MCPS)^19^ and Singapore’s SG10K Health project (SG10K)^20^. From the variant calling files provided by these projects, only single nucleotide variants (SNVs) meeting the “PASS” quality filter criteria were extracted. These variants were subsequently mapped to UniProt reference sequences using Ensembl VEP 110^51^, aligning with release versions 2023_03 for MCPS and SG10K, and 2024_02 for gnomAD v4.1.

### Genetic Ancestry Groups

To assign ancestry labels to samples, sequencing studies usually adopt statistical methods such as principal component analysis (PCA) to cluster genetically close samples and then train a classifier using the principal components as features on a set of reference samples with provided genetic ancestry labels. Although this approach and the use of genetic ancestry groups has important limitations^52^, comparing their representation in datasets and their interpretation by tools can provide a useful proxy for assessing the impact of ancestry bias in genetic studies.

In this study, we utilised variants associated with the following 14 ancestry groups as defined by the three population sequencing datasets:

- **gnomAD v4.1**: African/African American (AFR), Admixed American (AMR), Ashkenazi Jewish (ASJ), East Asian (EAS), Finnish (FIN), Middle Eastern (MID), non-Finnish European (NFE), and South Asian (SAS). The Amish (AMI) were excluded due to low sample count, whereas the Remaining Individuals (RMI) were filtered out to avoid ambiguity. The International Genome Sample Resource (IGSR)^53^, primarily based on the 1000 Genomes Project^54^, was utilised by the dataset as the reference to assign ancestry labels to the samples.
- **MCPS**: African (AMX), European (EMX), and Indigenous Mexican (IMX). The samples were assigned ancestry labels using the African (Yoruba) and European (Iberian) samples from the 1000 Genomes Project, and Indigenous Mexican samples from the Metabolic Analysis of an Indigenous Sample (MAIS) dataset^55^ as reference.
- **SG10K**: Chinese (CHI), Indian (IND), and Malay (MAL). The dataset used a reference panel consisting of 100 individuals from each of the three ancestry groups to assign ancestry labels to the samples.

### Variant Filtering

To ensure fair comparisons between ancestry groups, the analysis was restricted to single nucleotide missense variants with allele frequencies between 0.1% and 1%. This approach allowed the selection of variants with comparable penetrance across ancestry groups while avoiding biases from more extensively sequenced groups, which might otherwise over-represent rare variants and complicate comparisons. As a result, the number of variants analysed across ancestry groups and their allele frequency distributions were fairly consistent (Figure 1A).

### ROC AUC Calculations

The *metrics*.*roc_curve* function from the Scikit-learn Python package was used to compute the receiver operating characteristic (ROC) area under the curve (AUC) scores for each VEP using 64,066 variants labelled as (likely) benign and 51,575 variants labeled as (likely) pathogenic in ClinVar^22^ (downloaded on August 6, 2024). Pathogenic variants were treated as true positives, while benign variants were considered true negatives. For VEPs using an inverted scoring scale (where lower scores indicate greater predicted pathogenicity), the labels were inverted accordingly to maintain consistency.

### Variant Effect Predictors

A total of 90 VEPs were initially considered, with their scores obtained using the in-house pipeline described in Livesey et al.^6^. 25 VEPs were excluded since they had less than 50% coverage of variants in one or more ancestry groups. Additionally, to filter for VEPs with reasonably good performance, only VEPs with a ROC AUC score of >0.8 were retained, further filtering out 13 predictors (Figure S1). Applying these criteria resulted in the selection of 52 VEPs. Both Maverick^56^ and MOI-Pred^57^ provide in dominant and recessive variant predictions; for these, we ignored inheritance and used only the benign *vs* pathogenic scores.

### Rank-normalised Scores

To ensure consistency in comparisons, VEP scores for variants from all ancestry groups were collectively rank-normalised. This was achieved by firstly ranking the variant scores using the *rank* function from the Pandas Python package^58^, with identical values assigned the *average* rank. The resulting ranked scores were then normalised to a range between 0 and 1. For VEPs with an inverted scoring scale, where lower scores signify higher predicted pathogenicity, the scores were also inverted to ensure that predicted benign variants were assigned values closer to 0, and predicted pathogenic variants closer to 1.

## ACKNOWLEDGEMENTS

This project was supported by funding to JAM from the European Research Council (ERC) under the European Union’s Horizon 2020 research and innovation programme (grant agreement No. 101001169), a Lister Institute Research Prize Fellowship and by the Medical Research Council (MRC) Human Genetics Unit core grant (MC_UU_00035/9). JN is supported by an NMRC Clinician Scientist Award.

This study made use of data generated as part of the Singapore National Precision Medicine program funded by the Industry Alignment Fund (Pre-Positioning) (IAF-PP: H17/01/a0/007). This study made use of data/samples collected in the following cohorts in Singapore: (1) The Health for Life in Singapore (HELIOS) study at the Lee Kong Chian School of Medicine, Nanyang Technological University, Singapore (supported by grants from a Strategic Initiative at Lee Kong Chian School of Medicine, the Singapore Ministry of Health under its Singapore Translational Research Investigator Award (NMRC/STaR/0028/2017) and the IAF-PP: H18/01/a0/016); (2) The Growing up in Singapore Towards Healthy Outcomes (GUSTO) study, which is jointly hosted by the National University Hospital (NUH), KK Women’s and Children’s Hospital (KKH), the National University of Singapore (NUS) and the Singapore Institute for Clinical Sciences (SICS), Agency for Science Technology and Research (A*STAR) (supported by the Singapore National Research Foundation under its Translational and Clinical Research (TCR) Flagship Programme and administered by the Singapore Ministry of Health’s National Medical Research Council (NMRC), Singapore - NMRC/TCR/004-NUS/2008); (3) The Singapore Epidemiology of Eye Diseases (SEED) cohort at Singapore Eye Research Institute (SERI) (supported by NMRC/CIRG/1417/2015; NMRC/CIRG/1488/2018; NMRC/OFLCG/004/2018); (4) The Multi-Ethnic Cohort (MEC) cohort (supported by NMRC grant 0838/2004; BMRC grant 03/1/27/18/216; 05/1/21/19/425; 11/1/21/19/678, Ministry of Health, Singapore, National University of Singapore and National University Health System, Singapore); (5) The SingHealth Duke-NUS Institute of Precision Medicine (PRISM) cohort (supported by NMRC/CG/M006/2017_NHCS; NMRC/STaR/0011/2012, NMRC/STaR/ 0026/2015, Lee Foundation and Tanoto Foundation); (6) The TTSH Personalised Medicine Normal Controls (TTSH) cohort funded (supported by NMRC/CG12AUG17 and CGAug16M012). The views expressed are those of the author(s) and are not necessarily those of the National Precision Medicine investigators, or institutional partners. We thank all investigators, staff members and study participants who made the National Precision Medicine Project possible.

## DATA AVAILABILITY

Complete datasets associated with the analyses in this study and the list of contributing authors to the SG10K_Health Consortium is available at https://osf.io/wz2sb.

## SUPPLEMENTARY FIGURES

**Figure S1.**
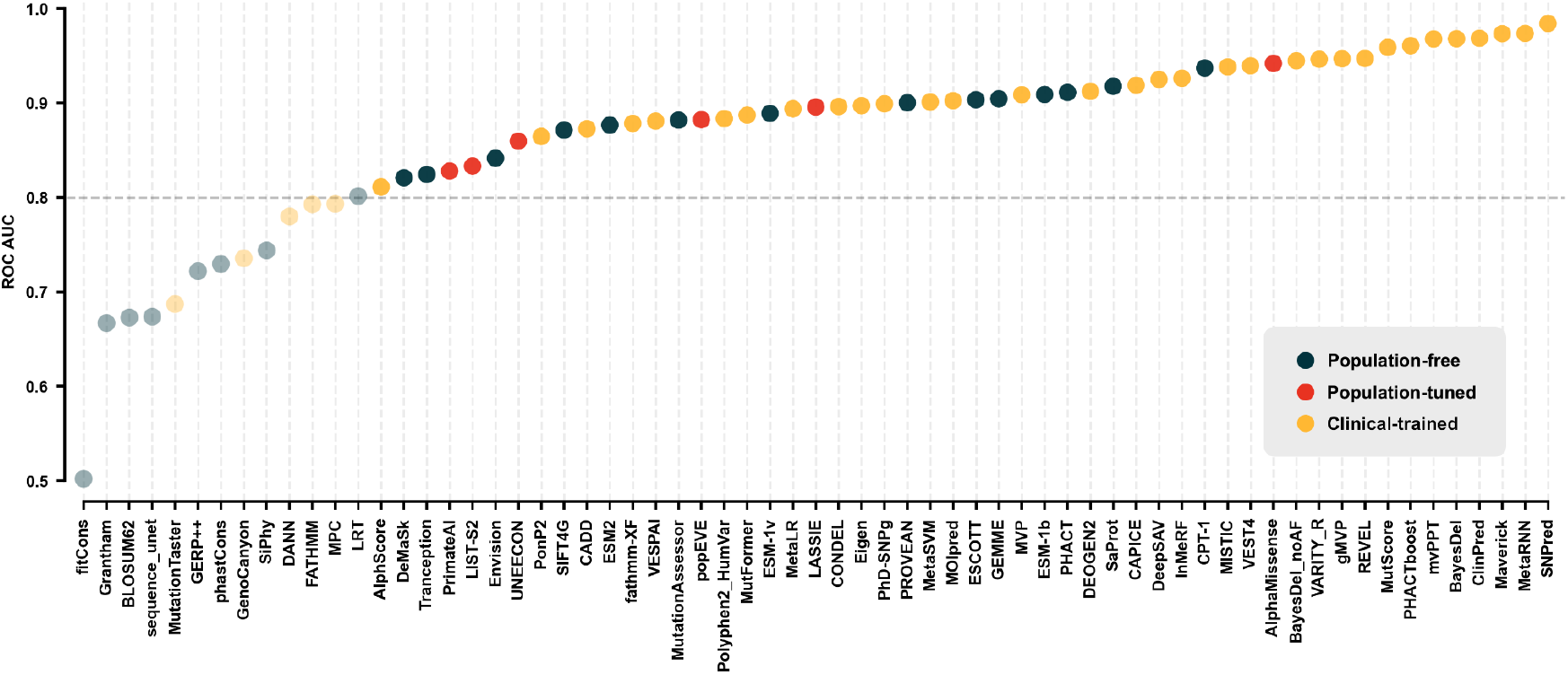
VEP performance in distinguishing known pathogenic and benign variants. The receiver operating characteristic (ROC) area under the curve (AUC) was calculated for each VEP to assess its ability to distinguish between missense variants classified as (likely) benign and (likely) pathogenic in ClinVar. VEPs were categorised into three classes based on their training methods: population-free, population-tuned, and clinical-trained. Only VEPs with a ROC AUC of >0.8 were considered in this study.

**Figure S2.**
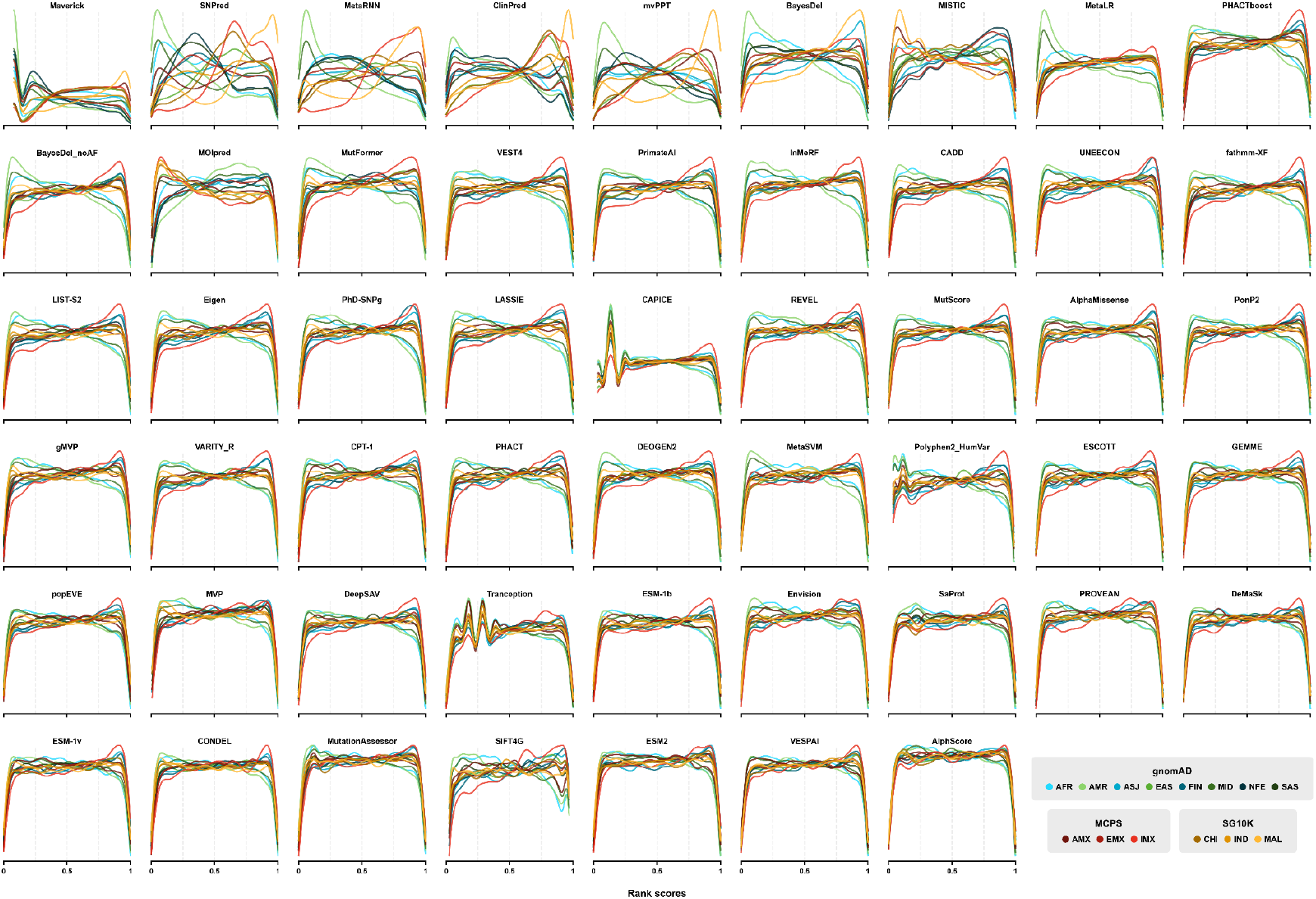
Score distribution of variants across ancestry groups. An extension of Figure 2A, displaying the rank score distributions of 52 VEPs – 14 population-free, 6 population-tuned and 32 clinical-trained models – across the 14 ancestry groups. The VEPs are ordered by decreasing standard deviation of mean rank scores.

**Figure S3.**
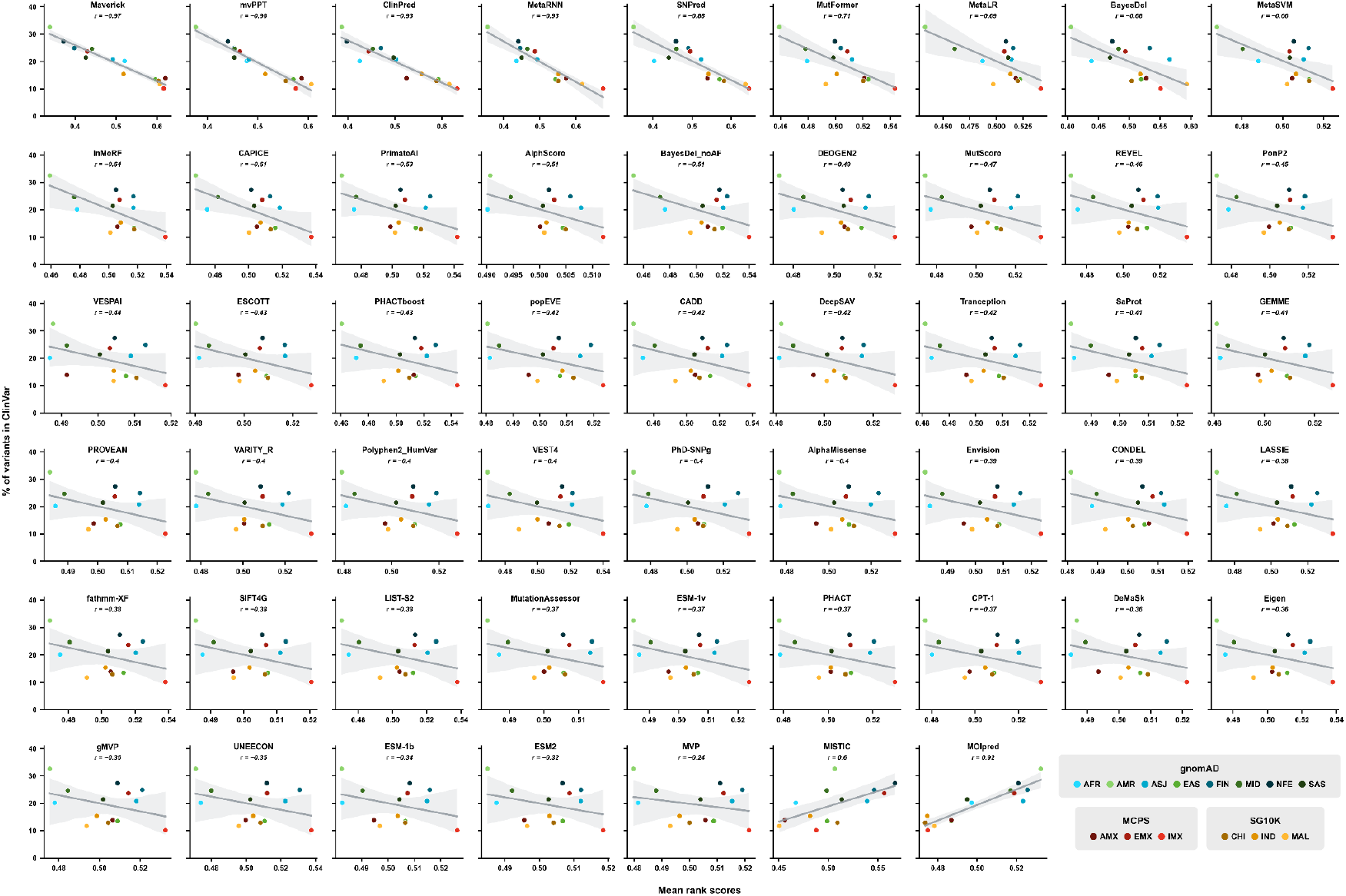
Correlation between mean rank scores and their representation in ClinVar across ancestry groups. An extension of Figure 2B, showing the correlation between the mean rank scores of 52 VEPs – 14 population-free, 6 population-tuned and 32 clinical-trained models – across the 14 ancestry groups and the percentage of these variants represented in ClinVar. The line indicates the best linear regression model fit, with the shaded region representing the 95% confidence interval. The Pearson correlation coefficient (*r*) quantifies the strength of this relationship, with VEPs ordered by increasing *r* values.

